# *Comparative virome analysis of individual shedding routes of* Miniopterus fuliginosus *bats inhabiting the Wavul Galge Cave, Sri Lanka*

**DOI:** 10.1101/2022.09.21.508883

**Authors:** Therese Muzeniek, Thejanee Perera, Sahan Siriwardana, Dilara Bas, Fatimanur Bayram, Mizgin Öruc, Beate Becker-Ziaja, Inoka Perera, Jagathpriya Weerasena, Shiroma Handunnetti, Franziska Schwarz, Gayani Premawansa, Sunil Premawansa, Wipula Yapa, Andreas Nitsche, Claudia Kohl

**Affiliations:** Robert Koch Institute, Centre for Biological Threats and Special Pathogens, Highly Pathogenic Viruses (ZBS 1), 13353 Berlin, Germany; Institute of Biochemistry, Molecular Biology and Biotechnology, University of Colombo, 00300 Colombo, Sri Lanka; IDEA (Identification of Emerging Agents) Laboratory, Department of Zoology and Environment Sciences, University of Colombo, 00300 Colombo, Sri Lanka; Robert Koch Institute, Centre for International Health Protection, Public Health Laboratory Support (ZIG 4), 13353 Berlin, Germany; Colombo North Teaching Hospital, 11010 Ragama, Sri Lanka

**Keywords:** metagenomic NGS, virome analysis, *Miniopterus fuliginosus*, Sri Lanka

## Abstract

Bats are described as the natural reservoir host for a wide range of viruses. Although an increasing number of bat-associated, potentially human pathogenic viruses were discovered in the past, the full picture of the bat viromes is not explored yet. In this study, the virome composition from *Miniopterus fuliginosus* bats inhabiting the Wavul Galge cave, Sri Lanka, was analyzed. To assess different possible shedding routes, oral swabs, feces and urine were collected and analyzed individually by using metagenomic NGS. The data obtained was further evaluated by using phylogenetic reconstructions.

Two different alphacoronavirus strains were detected in feces and urine samples. Furthermore, a paramyxovirus was detected in urine samples. Sequences related to *Picornaviridae, Iflaviridae*, unclassified *Riboviria* and *Astroviridae* were identified in feces samples, and further sequences related to *Astroviridae* in urine samples. No further viruses were detected in oral swab samples.

The comparative virome analysis in this study revealed a diversity in the virome composition between the collected sample types which also represent different potential shedding routes for the detected viruses. At the same time, several viruses were detected for the first time in bats in Sri Lanka.

The detection of two different coronaviruses in the samples indicates the potential general persistence of this virus species in *M. fuliginosus* bats. Based on phylogenetics, the identified viruses are closer related to bat-associated viruses with comparably low human pathogenic potential. In further studies, the seasonal variation of the virome will be analyzed to identify possible shedding patterns for particular viruses.

## Introduction

Bats are species-rich and taxonomically diverse mammals in the order *Chiroptera* that are distributed worldwide [1]. They represent a large group (20 %) of mammals and share unique features like their ability to fly. A number of viruses from different viral families, including human pathogenic viruses like Hendra and Nipah virus, coronaviruses, lyssaviruses and others have been associated with bats [2]. It is assumed that these viruses evolved together with their natural reservoir hosts. Because of this co-speciation and adaptation process, the viruses are often less pathogenic for their bat hosts. It is assumed that the bats’ immune system is adapted to and hence able to control virus infections without developing visible symptomatic diseases [3–5].

With the increasing focus on bat virus research, the detection of potentially zoonotic viruses with species-specific or family-specific PCR assays (e.g. paramyxoviruses, lyssaviruses, coronaviruses) has been a convenient standard method. However, this may have led to a bias focusing on certain viruses of particular interest [6–9]. To eliminate this disadvantage, metagenomic NGS methods (mNGS) for virus discovery were applied allowing for an untargeted and unbiased sequencing of novel viruses. The analysis of the whole virus composition in bat samples (viromes) allows to reduce the bias and to constantly increase the number of viral sequences deposited in sequence databases such as GenBank of the National Center for Biotechnology Information (NCBI) [10–12].

However, in several regions of the world the investigation of bats in their role as potential reservoir host of zoonotic viruses has been barely conducted [6]. In Sri Lanka, zoology is an important research field and ecological aspects of bats are well investigated [13]. They significantly contribute to the biodiversity and account for about a third of the Sri Lankan mammals with 30 different species [14]. Furthermore, they are essential for the maintenance of the ecosystem by providing ecoservices such as pollination, seed dispersal and insect control [13].

In contrast, only few studies have focused on bats as reservoir host for pathogens in Sri Lanka [15–18]. Here we present the first virome analysis of *M. fuliginosus* bats inhabiting Wavul Galge cave (Koslanda, Sri Lanka) in the interior of Sri Lanka. *M. fuliginosus* bats are roosting sympatrically with the other bat species *Hipposideros lankadiva, Hipposideros speoris, Rhinolophus rouxii* and *Rousettus leschenaultii*. In three individual field studies at different time points, we captured bats of all representative species and collected different sample material depending on availability [19]. Selected sample sets had been analyzed in previous investigations focusing on different research questions [17, 18, 20, 21]. In this study, we focus on the virome analysis of urine swabs (US), oral swabs (OS) and feces (F) collected from *M. fuliginosus* bats at one sampling point (July 2018). The presented results give insights into the virome composition of this bat species in the Wavul Galge cave and in general. Furthermore, the results reveal differences in viral shedding routes by analyzing the different sample types.

## Methods

The study was approved by the local government authority (Department of Wildlife Conservation, Sri Lanka, permit No. WL/3/2/05/18, issued on 10 January 2018). Catching and sampling of bats was carried out according to relevant guidelines and regulations of the Fauna and Flora Protection Ordinance [19].

### Bat Sampling

Sampling of bats inhabiting the Wavul Galge cave (Sri Lanka) was performed in March and July 2018 and January 2019 as described before [17]. For the results presented in this study, only bat samples from the species *M. fuliginosus* collected in July 2018 were included. 188 individuals of this bat species were captured and sampled with adequate personal protective equipment. Different sample types were collected depending on availability. From *M. fuliginosus* a total of 187 oral swabs (OS), 102 urine swabs (US) and 77 fecal pellets (F) were taken and analyzed.

The general workflow of subsequent laboratory work and bioinformatic analysis of NGS data is shown in Figure 1.

**Figure 1:**
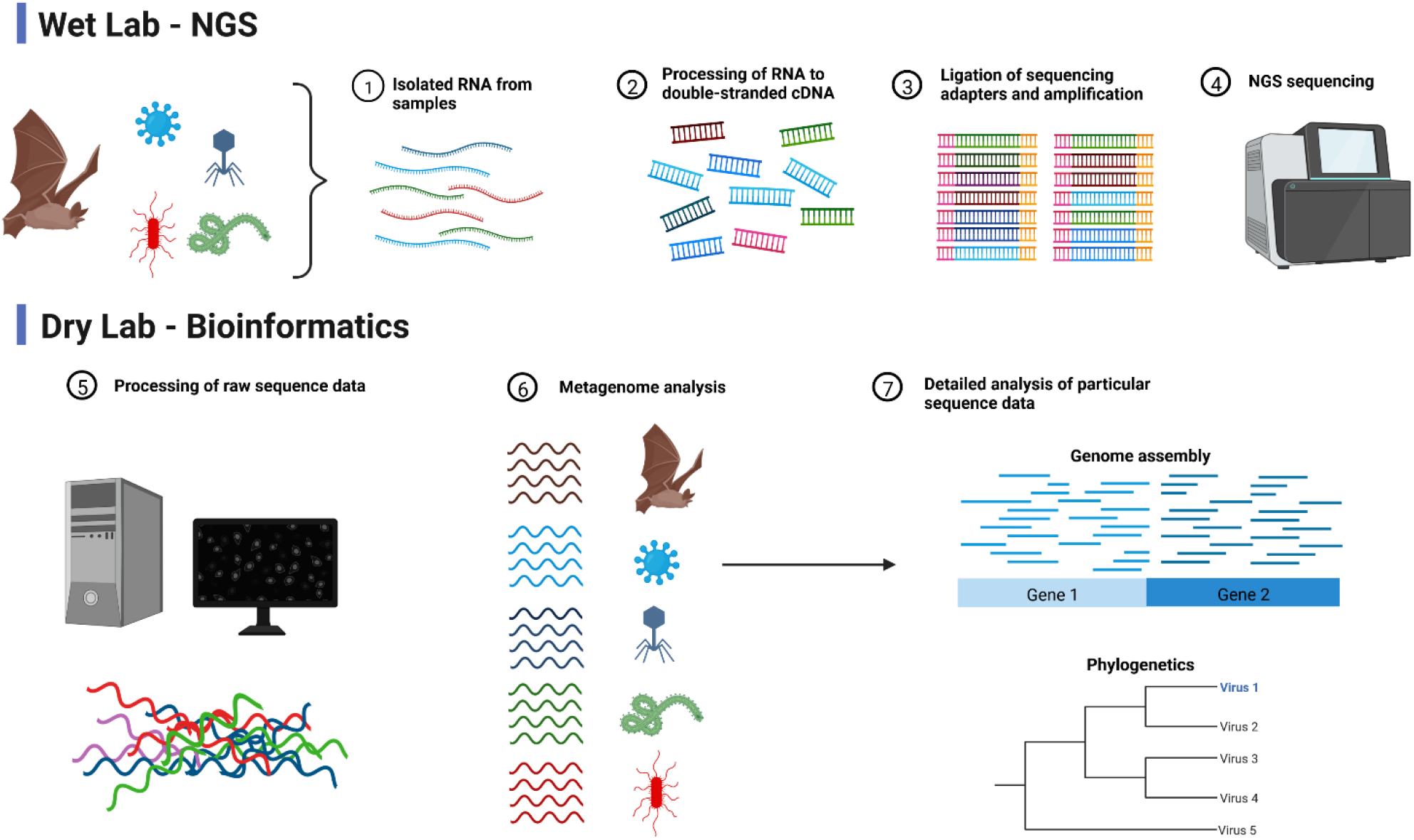
Schematic illustration of the general NGS workflow (wet lab) with subsequent bioinformatic analysis of obtained data (dry lab). The goal is to analyze the general virome composition (6) of the respective sample and to further characterize novel viruses based on the obtained sequencing data (7). Created with BioRender.com

### Metagenomic NGS

The processing of samples was conducted with appropriate precautions in biosafety-2 laboratories. All samples were initially processed by adding 500 µL of sterile PBS. For oral swabs and urine swabs, samples were mixed by vortexing before 140 µL were used for extraction with the Viral RNA Mini Kit (QIAGEN, Hilden, Germany). Fecal pellets were homogenized with sterile ceramic beads by using the FastPrep-24 device (MP Biomedicals, Eschwege, Germany), followed by centrifugation and extraction of 140 µL of the supernatant.

Prior to further processing a maximum of 10 RNA extracts were pooled by taking 5 µL per sample to obtain a final volume of 50 µL. Pools were prepared per sample type, resulting in 18 OS pools, 13 US pools and 8 F pools. The pooled RNA samples were treated with DNase at 37 °C by using a TURBO DNA-free Kit (Invitrogen, Carlsbad, CA, USA), followed by RNA transcription into cDNA by using the SuperScript IV Reverse Transcriptase reagent (Invitrogen) and random hexamer primers. Second strand was synthesized by using the NEBNext® Ultra™ II Non-Directional RNA Second Strand Synthesis Module (New England Biolabs, Ipswich, MA, USA) according to the manufacturer’s instructions. dsDNA was purified by using the Agencourt AMPure XP bead system (Beckman Coulter Life Sciences, Krefeld, Germany) and the final DNA concentration was determined by using a NanoDrop™ 1000 Spectrophotometer (Thermo Fisher Scientific, Hennigsdorf, Germany). Sample libraries were sequenced on a HiSeq 2500 or a NextSeq 550 sequencing device (Illumina, San Diego, CA, USA) with a paired end read output of 2 × 250 bp (HiSeq) or 2 × 150 bp (NextSeq) and a total output of up to 8 million reads per pool. A detailed overview of the pools, included samples and read output is given in the supplementary material (Table ST1).

### NGS data analysis and virome assembly

Prior to the NGS data analysis a quality trimming of sequencing reads was performed by using the tool Trimmomatic v0.39 [22]. For further analysis the diamond alignment tool was utilized to compare trimmed reads to the non-redundant protein virus database (NCBI, RefSeq release 210 from 3 January 2022) by using the BLASTx algorithm [12, 23]. The results were analyzed and the distribution of assigned viral reads per pool was evaluated in MEGAN [24]. This was also compared between the different sample types and among the pools of each sample type. Selected results were subsequently blasted to the whole NCBI database by using the BLASTn algorithm. For a number of hits the full genome sequences were downloaded and used as a reference for mapping the trimmed NGS data to identify more reads and to evaluate the mapping quality visually in Geneious Prime software (version 2020.2.3, Biomatters Ltd., Auckland, New Zealand). Wherever possible, the nucleotide identities to related strains were calculated for the longest contig assembled on a conserved gene such as the RNA polymerase gene.

For viral sequences of high interest, suitable primers were designed on contigs supported by high read coverage. Subsequently, the presence of the novel viral sequences was confirmed in the initial sample pools by using conventional PCR under standard conditions (available on request). PCR products appearing as bands in the analytic agarose gel were purified and Sanger sequenced when sufficient quantity was reached.

### Phylogenetic reconstruction

For selected viral sequences of interest, suitable contigs were used for phylogenetic reconstruction. For this purpose, a number of reference sequences were downloaded from NCBI database and a gap-free nucleotide alignment was calculated by using the MAFFT algorithm [25]. The phylogenetic trees were calculated based on this alignment by using the Bayesian MCMC algorithm (MrBayes version 3.2.6) [26]. A suitable model for each alignment was estimated by using the Akaike information criterion (AIC) prediction with the tool JModelTest [27, 28]. The calculation parameters were different depending on the virus and are specified in the results section (see figure legends Figure 4, Figure 5 and Figure 6), respectively.

## Results

Following mNGS, the sequence reads were trimmed and analyzed separately per pool as described in the methods section. Comparison of the pools per sample type revealed mainly homogenous distributions of viral reads assigned to the respective viral orders (see supplemental Figures SF1–SF3). For further analysis, viral reads of all pools per sample type were combined and the assigned viral reads were compared between OS, F and US by using MEGAN as shown in Figure 2.

**Figure 2.**
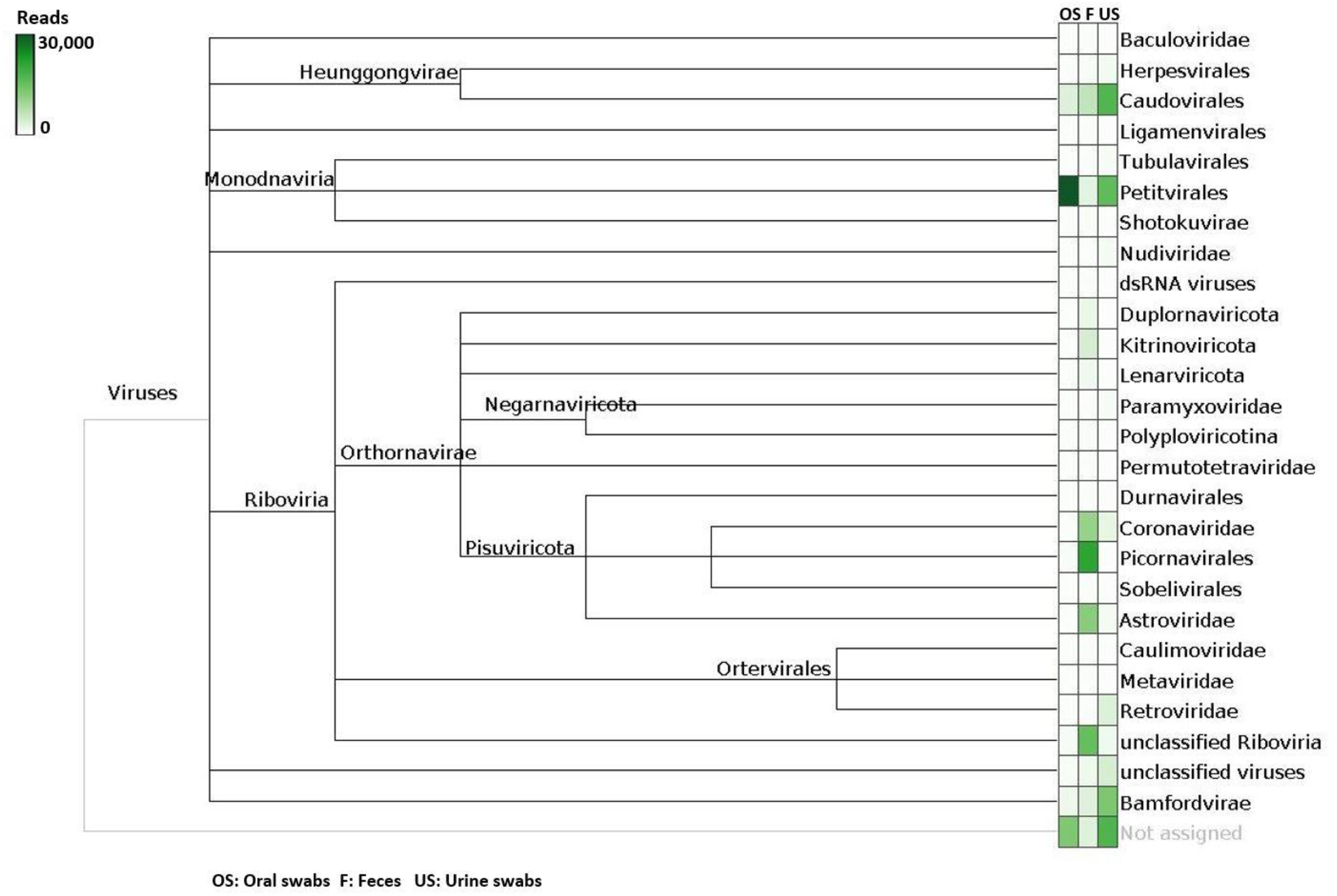
Normalized comparison of viral hits from different sample types in MEGAN after diamond BLASTx. Sample types are depicted in the following order: oral swabs (OS), feces (F), urine swabs (US). The intensity of green color represents the quantity of reads assigned to the respective viral family or order.

As shown in Figure 2, bacteriophages (e.g. *Caudovirales, Petitvirales*) were found in all sample types but in different amounts. For OS, no other viral hits of interest were assigned from the NGS data. Viral reads belonging to six different families (*Coronaviridae, Iflaviridae, Picornaviridae, Astroviridae, Paramyxoviridae* and unclassified *Riboviria*) were found in F samples and confirmed in silico and also via PCR. In the US samples, viral reads of three families (*Paramyxoviridae, Coronaviridae* and *Astroviridae*) were detected and confirmed in silico and also via PCR.

Some of the other preliminary results from BLASTx (depicted in Figure 2) could not be confirmed in the subsequent in silico analysis and were considered as possible cross-contaminations from the NGS run (e.g. *Herpesvirales* and *Bamfordvirae*) as commonly described [29]. An overview of the confirmed results after quality checks is given in Table 1, specifying details of the obtained NGS data, related virus and novel identified virus strains.

**Table 1:**
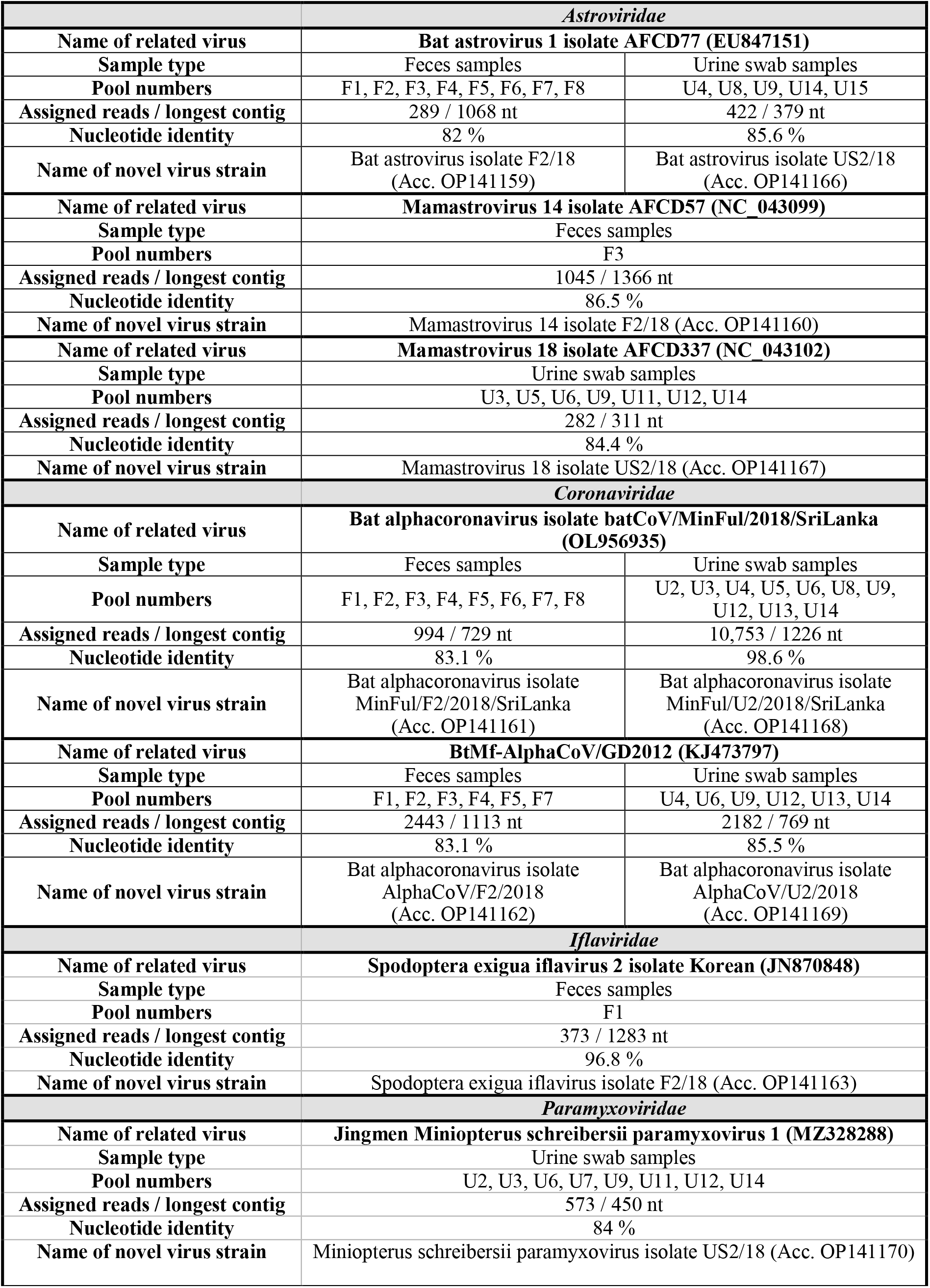
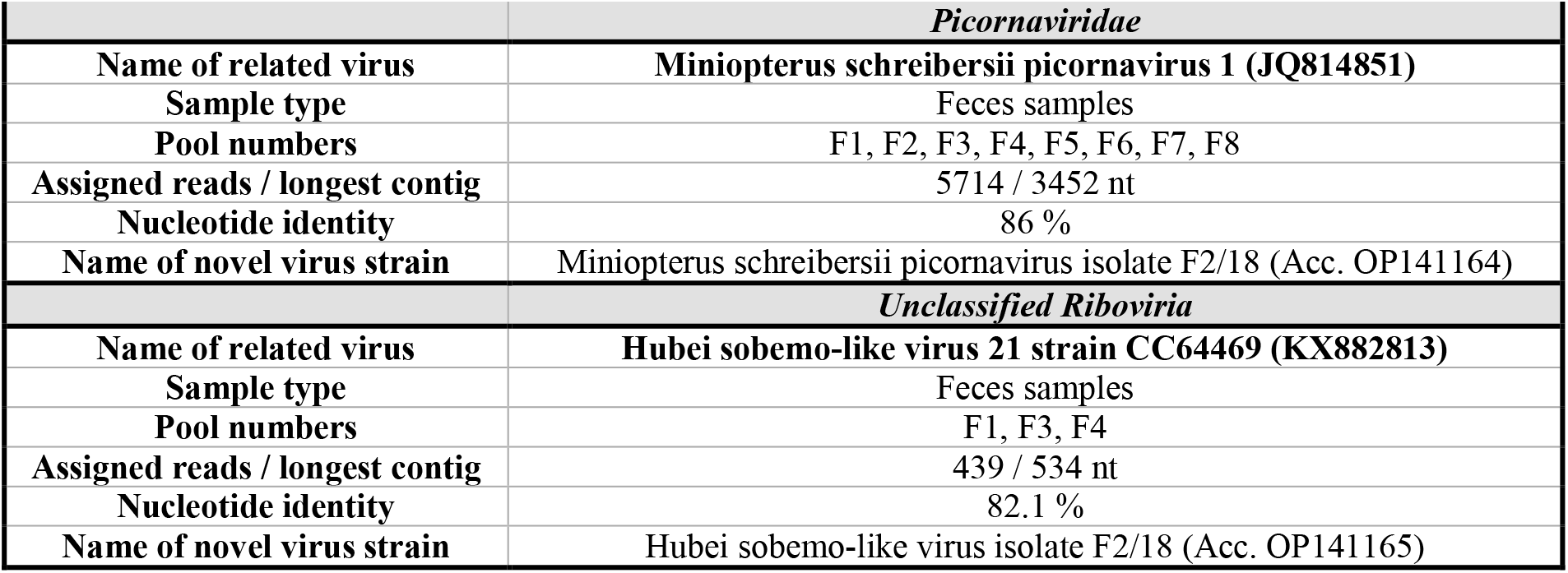
Overview of the results obtained from mNGS data analysis of feces and urine swab samples from *M. fuliginosus* bats collected in July 2018. The table indicates the related virus, sample types and pool numbers from which the results were obtained, including the number of assigned reads, longest contig and nucleotide identity. The name and accession number of the novel virus strain as uploaded to GenBank are indicated, respectively.

For the families *Coronaviridae, Picornaviridae, Astroviridae* and *Paramyxoviridae*, the presence of viral sequences was furthermore confirmed in the original RNA pools (see Table 1) by using PCR and specifically designed primer based on the NGS data. The remaining viruses (*Iflaviridae* and unclassified *Riboviria*) were solely confirmed with in silico analysis of the data.

From all viral assemblies, either the longest contig or the contig used for phylogenetic reconstruction was uploaded to GenBank. These sequences of newly detected virus strains were named in relation to the closest related reference virus and with respect to current ICTV classification criteria (see Table 1).

### Astroviridae

Astroviruses (n=3) were found in F and US samples. In F samples, 1045 reads were assigned to Mamastrovirus 14 isolate AFCD57 (NC_043099) with a nucleotide identity of 86.8 % on the longest contig (1366 nt), located overlapping on the polyprotein 1AB and capsid protein CDS. Further 289 reads from F samples were assembled to Bat astrovirus 1 isolate AFCD77 polyprotein 1AB gene (EU847151) with a nucleotide identity of 82 % on the longest contig (1068 nt).

A total of 422 reads from US samples were also assembled to EU847151, with a nucleotide identity of 85.6 % on the longest contig (379 nt).

From US samples further 282 reads were assembled to Mamastrovirus 18 isolate AFCD337 (NC_043102) with a nucleotide identity of 84.4 % on the longest contig (311 nt), located on the polyprotein 1AB gene.

Because of the lack of overlapping sequences, phylogenetic reconstruction was calculated exemplarily with Bat astrovirus isolate F2/18 (OP141159); the results are shown in Figure 3.

**Figure 3:**
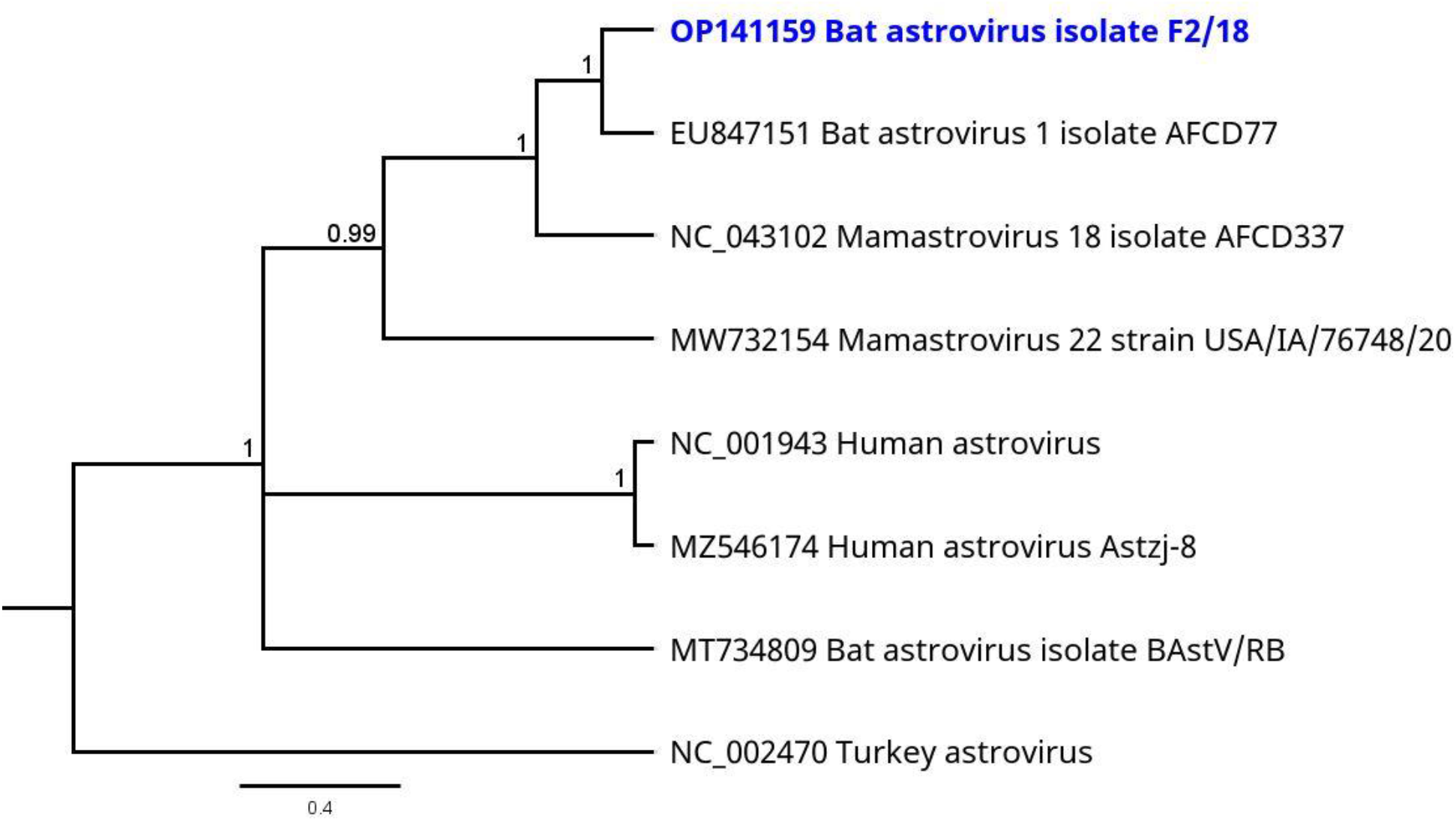
Phylogenetic tree based on the mNGS sequences obtained from the virome of F sample pools (highlighted in blue) and a selection of other astrovirus species. Turkey astrovirus (NC_002470) was used as outgroup. Phylogenetic reconstruction was calculated with the Bayesian MCMC algorithm: 500,000 generations were calculated with a subsampling frequency of 100 and a burn-in of 10 %. The substitution model GTR was selected with a gamma-distributed rate variation. Visualized as molecular clock with uniform branch lengths.

### Coronaviridae

Two coronaviruses (CoV) were found in US and F samples, respectively. A total of 994 reads from F samples and 10,753 reads from US samples were mapped to the bat alphacoronavirus isolate batCoV/MinFul/2018/SriLanka (OL956935) with nt identities on conserved ORF1B CDS of 80.36 % (F, 668 nt contig) and 98.37 % (US, 735 nt contig). In addition, 2443 reads from F sample pools and 2182 reads from US sample pools were mapped to BtMf-AlphaCoV/GD2012 (KJ473797), an alphacoronavirus HKU8 strain from China. Nucleotide identities on the conserved ORF1B CDS were calculated with 82.9 % (F, 1113 nt contig) and 85.5 % (US, 769 nt contig). For phylogenetic reconstruction, contigs on the ORF1B CDS were selected, respectively. Because of the lack of overlapping contigs, phylogeny was calculated separately for the sequences from F and US samples but by using the same reference strains including common human pathogenic CoVs. For the conserved ORF1B CDS, overlapping contigs of 182 nt (F samples) and 224 nt (US samples) were obtained. Figure 4 shows the phylogenetic reconstruction of CoV from US sample pools.

**Figure 4:**
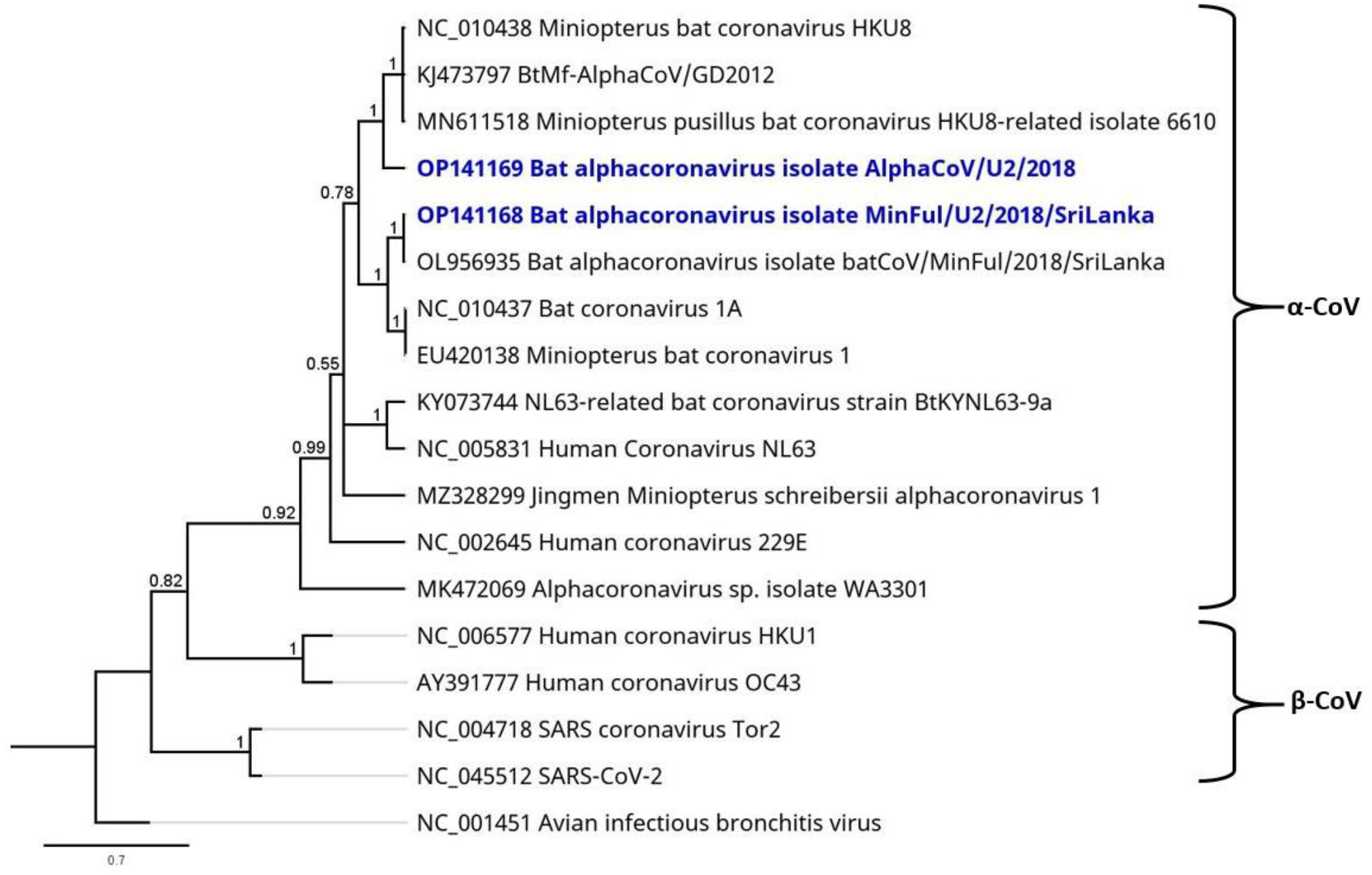
Phylogenetic tree based on the mNGS sequences obtained from the virome of US sample pools (highlighted in blue) and a selection of α-CoVs and β-CoVs as specified. For use as outgroup, the γ CoV avian infectious bronchitis virus (NC_001451) was also included in the calculation. Phylogenetic reconstruction was calculated with the Bayesian MCMC algorithm: 1,000,000 generations were calculated with a subsampling frequency of 100 and a burn-in of 10 %. The substitution model GTR was selected with a gamma-distributed rate variation. Visualized as molecular clock with uniform branch lengths.

The calculation confirms the presence of two different strains that are allocated to different branches inside the genus *Minunacovirus*. The identified Bat alphacoronavirus batCoV/MinFul/2018/SriLanka US2/2018 clusters with strains of the species Miniopterus bat coronavirus 1, whereas the other Miniopterus AlphaCoV strain US2/2018 clusters with HKU8-related strains.

### Iflaviridae

An iflavirus was found in F samples with a total of 373 reads assembled to Spodoptera exigua iflavirus 2 isolate Korean (JN870848), covering 93.6 % of the genome. The longest contig of 1283 nt, located at the beginning of the polyprotein CDS, shares a nucleotide identity of 96.8 % to the related Spodoptera exigua iflavirus 2.

### Paramyxoviridae

A paramyxovirus (PMV) was found in US samples and confirmed by PCR (compare Table 1). A total of 573 reads were mapped to the full genome of Jingmen Miniopterus schreibersii paramyxovirus 1 (MZ328288). The longest contig (450 nt) on the conserved L gene showed highest nucleotide identities to this strain (84 %) and to the partial genome of Miniopterus schreibersii paramyxovirus isolate Bat Ms-ParaV/Anhui2011 (KC154054; 84.6 % nt identity). This contig was also selected for phylogenetic reconstruction. The phylogenetic tree of 16 paramyxoviruses including the novel sequence from Sri Lanka and selected human pathogenic strains is shown in Figure 5.

**Figure 5:**
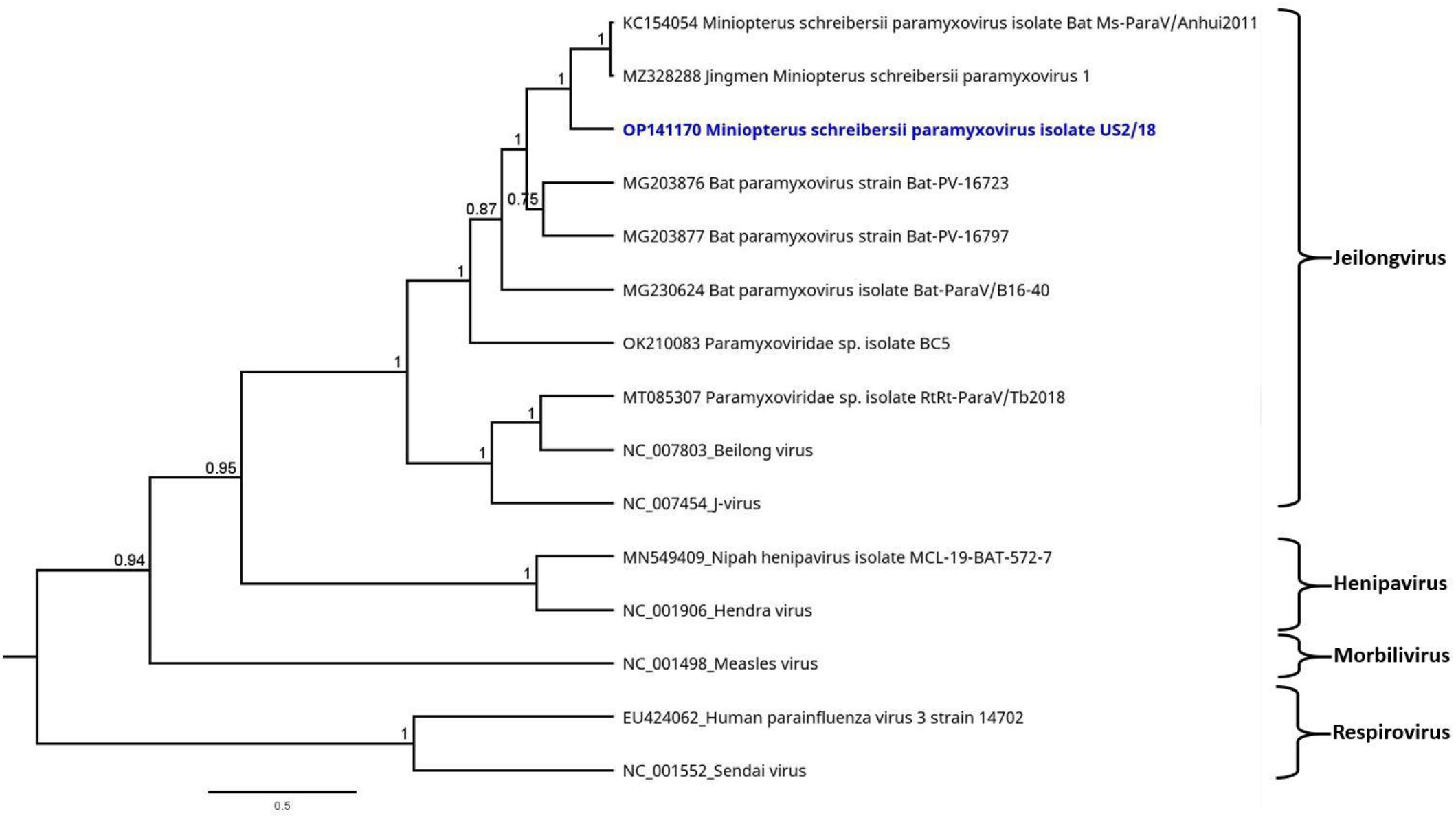
Phylogenetic tree based on the mNGS sequences obtained from the virome of US sample pools (highlighted in blue) and a selection of PMVs. Sendai virus (NC_001552) was selected as outgroup for the calculation. Phylogenetic reconstruction was calculated with the Bayesian MCMC algorithm: 1,000,000 generations were calculated with a subsampling frequency of 100 and a burn-in of 10 %. The substitution model GTR was selected with a gamma-distributed rate variation. Visualized as molecular clock with uniform branch lengths

The phylogenetic reconstruction confirms that the novel paramyxovirus from Sri Lanka is closely related to the two PMV strains as described before. Other Miniopterus-related PMVs from China cluster in the same branch of the tree, representing the subgenus Jeilong virus. Apart from this, the subgenera Henipavirus, Morbillivirus and Respirovirus were allocated to distinct branches of the tree, respectively. These subgenera also include the human pathogenic strains that were selected for this phylogenetic reconstruction.

### Picornaviridae

A Picornavirus was found in F samples, a total of 5714 NGS reads were assembled to Miniopterus schreibersii picornavirus 1 (JQ814851), covering most parts of the genome. The longest contig of 3452 nt, located at the end of the polyprotein CDS, shares a nucleotide identity of 86 % to the Miniopterus schreibersii picornavirus 1. For phylogenetic reconstruction, a well-covered contig of 311 nt was selected on the highly conserved 2C peptide on the picornaviral polyprotein. The phylogenetic reconstruction is illustrated in Figure 6.

**Figure 6:**
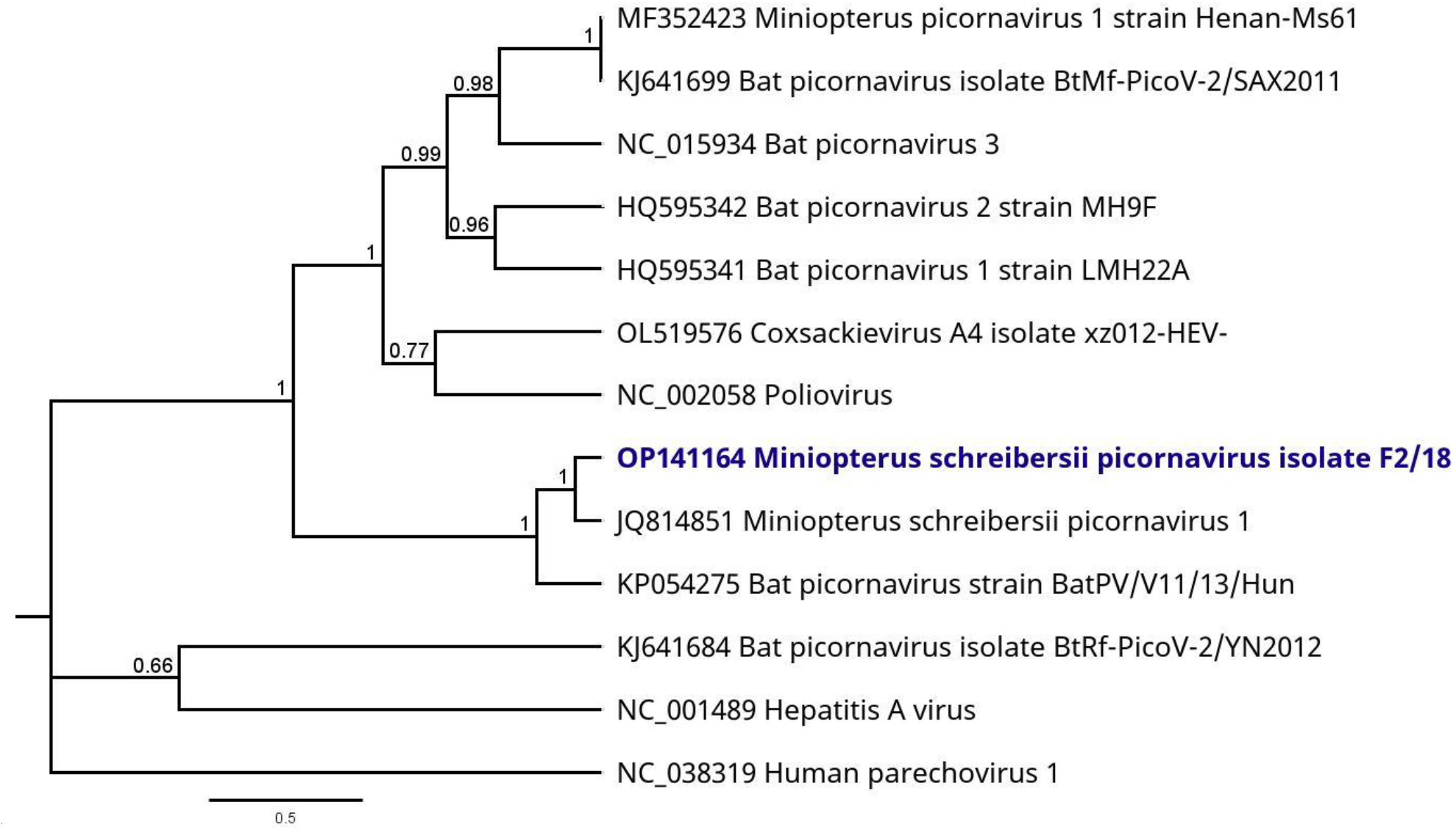
Phylogenetic tree based on the mNGS sequences obtained from the virome of F sample pools (highlighted in blue) and a selection of picornaviruses. Human parechovirus 1 (NC_038319) was selected as outgroup for the calculation. Phylogenetic reconstruction was calculated with the Bayesian MCMC algorithm: 1,000,000 generations were calculated with a subsampling frequency of 100 and a burn-in of 10 %. The substitution model GTR was selected with a gamma-distributed rate variation. Visualized as molecular clock with uniform branch lengths.

As shown in Figure 6, the novel bat picornavirus strain from F samples clusters monophyletically with its closest related strain Miniopterus schreibersii picornavirus 1 and the bat picornavirus strain BatPV/V11/13/Hun, both obtained from *Miniopterus schreibersii* bats. Other bat- and Miniopterus-hosted picornaviruses were allocated to different branches of the phylogenetic tree. The human pathogenic representative picornaviruses are clearly distant from the novel Sri Lankan strain.

### Unclassified Riboviria

From F samples, 439 reads were assembled to a Hubei sobemo-like virus 21 strain CC64469 (KX882813) with a nucleotide identity of 82.1 % on the longest contig (534 nt).

### Phages

A number of phages were found in each sample type as described. Considerable amounts of reads were assigned to the bacteriophage orders *Caudovirales* (OS: 576; F: 1278; US: 663,683) and *Petitvirales* (OS: 44,741; F: 235; US: 554,119), respectively.

## Discussion

In our study we analyzed the virome of *M. fuliginosus* bats inhabiting the Wavul Galge cave in the interior of the island of Sri Lanka. By taking orals swabs, urine swabs and feces, we aimed to examine if different viruses are shed by distinct shedding routes represented by the individual sample types. This assumption was confirmed and we have been able to obtain different virome compositions for each sample type.

In oral swab samples, the primary viral hits were assigned to phages belonging to the orders *Caudovirales* and *Petitvirales*. The presence of any other viral sequences in oral swab samples was not confirmed after quality assessment of analyzed NGS data. However, the detection of *Caudovirales* and *Petitvirales* in OS serves as proof of principle also for this sample type.

Apart from this, different other viruses were detected in the feces and urine swab samples as discussed in the following sections.

### Phages and viruses from other presumed reservoir hosts

Sequence reads assigned to phages were found in high numbers in all three sample types. Although these were not inspected in detail in this study, the large number and variety of reads also indicates the presence of numerous bacteria in the collected samples. In combination with 16S bacterial metagenomic analysis, these results will be interesting to be further explored in future studies.

In feces samples, viral sequences were identified matching Hubei sobemo-like virus 21 strain CC64469 (China), belonging to unclassified *Riboviria*. This virus was originally isolated from invertebrates within the phylum *Annelida* [30]. Most probably a worm carrying this virus was taken in as nutrition by the bat, passaged and excreted with the feces afterwards.

In addition to this, viral sequences assigned to Spodoptera exigua iflavirus 2 (Korea) were identified in feces samples. This virus belonging to the family *Iflaviridae* within the order *Picornavirales* was isolated from insects of the genus *Spodoptera* (moths) [31]. Here again it is very likely that the virus was taken in with moths for nutrition, passaged and excreted afterwards. In both examples, it is impossible to conclude whether the virus infected the bats as well or was merely digested, passaged and excreted. For this, tissues and organs would need to be investigated.

However, the identified phages as well as these two examples show that the viromes derived from NGS data are complex and do not only comprise the bat-related viruses. Moreover, the data also reveal a number of other viruses derived from the inherent bacterial flora within the bat as well as viruses derived from insects serving as nutrition for the bats. This in turn has the potential to analyze virome compositions of whole habitats, including different organisms. This may be the basis or part of further investigations regarding the bacterial flora and dietary habits of the bats.

### Astroviridae

The viruses within the family *Astroviridae* are common in a wide range of birds (genus *Avastroviruses*) and mammals (genus *Mamastrovirus*), including bats and humans [32]. Human astrovirus infections mainly cause gastroenteritis in children [33]. Members of this family have a high genetic diversity depending on the respective host species and a wide host range (birds, mammals). However, the zoonotic potential of bat-hosted astroviruses is widely unexplored as yet [34]. Members of the mamastrovirus genus are classified based on the amino acid sequences of the capsid region and can be further divided into species depending on the host and other genetic criteria [35]. Based on the analyzed sequence data, the examined bats carried multiple astrovirus strains. With this, bat astrovirus sequences were detected for the first time in a bat species from Sri Lanka, namely *M. fuliginosus*.

From US samples, Bat astrovirus isolate US2/18 (OP141166) and Mamastrovirus 18 isolate US2/18 (OP141167) were identified, whereas from F samples Bat astrovirus isolate F2/18 (OP141159) and Mamastrovirus 14 isolate F2/18 (OP141160) were identified. All three viruses were originally detected in *Miniopterus* bat species, which may indicate a high host specificity of these viruses. In addition, the circulation of different bat astrovirus strains within one bat population is conceivable, as already reported in other studies [36–38].

Phylogenetic reconstruction allowed for a rough classification of the Bat astrovirus isolate F2/18 from Sri Lanka, as an example. Although only few suitable reference strains were available in the databases, the phylogeny distinguished astroviruses of bats from astroviruses of other vertebrates (turkey, human) by allocation to different tree branches.

Most available bat astroviral references from the databases contained partial genomes only, located at different regions. Further sequence analysis and phylogenetic comparison between all newly obtained astrovirus sequences from Sri Lanka were therefore not possible with the available data. Consequently, we cannot finally prove the presence of multiple astrovirus strains in the collected samples. It may be possible that the sequences are originally derived from a single astrovirus but were mapped to the different partial genomes obtained from the database. Therefore, virus isolation of these viruses followed by in-depth sequencing of the missing genome sequences could be helpful in order to obtain full genome data or at least full gene sequences. With further sequence information it could be examined whether the astroviral reads were actually derived from different strains or if they belonged to a single astrovirus within the samples.

### Coronaviridae

Presence of two different CoVs in *M. fuliginosus* bats was confirmed in F and US samples. Both related virus strains were originally detected in *Miniopterus* bats [20, 39, 40]. The CoV full genome from Sri Lanka (OL956935) was derived from rectal swabs collected during the same bat sampling session as this study and reported previously [20]. With the virome sequence data of F and US samples in this study, we were able to confirm the presence of this virus strain in other sample types. In addition, we identified Bat alphacoronavirus isolate AlphaCoV/F2/2018 and Bat alphacoronavirus isolate AlphaCoV/U2/2018 which are closely related to Miniopterus bat coronavirus HKU8 strains. All identified viruses belong to *Minunacoviruses*, an α-CoV subgenus containing *Miniopterus*-hosted bats. The slight differences between the two virus species within the subgenus are also visible in the phylogenetic reconstruction of different CoVs. Representative strains of the virus species Miniopterus bat coronavirus 1 and Miniopterus bat coronavirus HKU8 were divided into two different clusters within the branch of *Minunacoviruses*. Co-existence of these two virus species and different CoVs in general have already been reported before [41–43]. Therefore, it can be assumed that these two or even more CoV strains circulate steadily in this population of *M. fuliginosus* bats. Regarding the zoonotic potential of the detected CoVs, further investigation of full genes and specific receptor-binding domains would be necessary for more precise statements. Based on the available data and phylogenies calculated from this, we did not find indications for a human pathogenic potential of the detected viruses [17, 20]. Human pathogenic CoVs that cause outbreaks and pandemics like SARS, MERS and Covid-19 all belong to the genus of β-CoV and are genetically diverse from the genus α-CoV. Although mildly human pathogenic viruses such as HCoVs NL63 and 229E are represented in the α-CoV genus as well, the phylogeny ranks these species as rather distantly related to the *Minunacoviruses*.

### Paramyxoviridae

Representatives of *Chiroptera*-hosted PMVs are able to cause zoonotic diseases in humans; therefore this virus family was of high interest for the virome analysis of the bat samples [44]. The detection of PMVs in US samples was expected, as this is the usual shedding route of these viruses [45]. The presence of PMVs in the collected US samples was also detected via semi-nested PCR [7] and Sanger sequencing as reported before [18]. These results indicated the presence of multiple PMVs in the bat population. The co-circulation of multiple PMVs is common in bats and has been reported before [46, 47]. With the obtained NGS data of this study we were able to confirm PMVs in the samples by using another molecular virus detection method. With this, a proof of principle of the methodical approach was possible. However, the NGS data did not reveal enough sequence information to confirm multiple PMV strains. For this purpose, NGS of single US samples instead of pools and comparison of sequence data between individual bats will be necessary.

As expected, the identified Miniopterus schreibersii paramyxovirus isolate US2/18 (OP141170) has the highest similarities to other *Miniopterus*-hosted PMVs. In the phylogenetic tree, all *Miniopterus*-derived strains are assigned to the group of Jeilong virus (Figure 5). The other branches of the phylogenetic tree depict only a small number of representative strains, including human pathogenic PMVs, whereas the actual *Orthoparamyxovirinae* subfamily is notably more diverse [44, 48]. In accordance to this, different PMVs can cause diseases of different severity in humans (e.g. Human parainfluenzavirus vs. Nipah virus), whereas other PMVs have a rather low zoonotic potential. The preliminary analysis of the limited sequence data and the phylogenetic reconstruction give no indication of a human pathogenic potential of the novel PMVs. Further sequence data will be needed to allow for a detailed and complex analysis and taxonomic classification of the novel PMV strain [49].

### Picornaviridae

The family of picornaviruses is a highly diverse family with 68 genera and 158 virus species according to ICTV [35]. They are globally distributed in a number of bat species including *Miniopterus* bats [50, 51]. Additionally, they are found in a number of other host species including birds, livestock and humans. Cross-species transmissions between different bats or mammals are possible as well as zoonotic transmissions to humans [52]. In humans, picornaviruses such as enterovirus, rhinovirus, coxsackievirus, hepatovirus A and human parechovirus can cause diseases of the nervous system and the respiratory and gastrointestinal tracts [53].

A number of sequences related to Miniopterus schreibersii picornavirus 1 were identified in feces samples, which is a described shedding route of bat picornaviruses [54]. The novel strain from Sri Lanka Miniopterus schreibersii picornavirus isolate F2/18 (OP141164) shares an identity of 86 % to the reference strain from China [55]. For phylogenetic reconstruction in this study, a suitable sequence on the 2C peptide was selected which is a highly conserved region on the picornaviral polyprotein and therefore suitable for this analysis [56]. The phylogenetic reconstruction included several bat picornaviruses and human pathogenic strains. The novel picornavirus strain from Sri Lankan *M. fuliginosus* bats was assigned to a branch with other *Miniopterus*-hosted picornaviruses, and the human pathogenic species were assigned to other branches of the tree. The available results did not indicate a human pathogenic potential of the identified picornavirus. Although the phylogenetic analysis was limited to a small proportion of the *Picornaviridae* family, we were able to get a general idea on phylogenetic relationships of the novel bat picornavirus from Sri Lanka. For proper species classification, a full protein sequence analysis of P1, 2C, 3C and 3D proteins will be necessary but was not possible from the obtained data. However, the results represent the first detection of a picornavirus in the bat species *M. fuliginosus* from Sri Lanka.

## Conclusion

We were able to analyze the different compositions of the *M. fuliginosus* virome by potential shedding route obtained from oral swabs, urine swabs and feces samples. Depending on the sample types, different viruses were detected via NGS analysis, corresponding to their typical shedding routes, respectively.

Independent of the sample type, we were able to detect the co-existence of astroviruses, coronaviruses, paramyxoviruses and picornaviruses circulating simultaneously in the

*M. fuliginosus* bat population. Co-existence of these viruses may be common in bats, and even a co-speciation of virus species with their specific host is discussed [57, 58]. It is assumed that also virus–virus interaction is possible and may influence the host, resulting in very specific viral shedding patterns depending on the virome composition [47]. In future, the epidemiological consequences of co-existing viruses in the bats should be further examined.

It is remarkable that mainly bacteriophages were identified in oral swab samples, although saliva is also known as common shedding route for other virus families. Lyssaviruses including rabies-related viruses would have been most likely to be found in this sample type. If these viruses are prevalent in the bats they are excreted by salivary glands and therefore shed with the saliva [59]. However, the excretion of viruses is generally affected by seasonal patterns and may have been low at the respective sampling point. Since we only used non-invasive sampling methods and could not examine bat brain or other tissue samples, we cannot conclude whether or not such viruses were prevalent in bat organs at the time of bat sampling. A long-term and frequent bat sampling will help to understand seasonality and shedding patterns of different viruses of interest. Probably the virome of all different sample types would change over time as it is influenced by seasons and environmental factors like temperature, humidity, rainfall, migration and reproduction cycles [1, 47, 60, 61].

In general, all collected sample types also represent possible transmission routes from bats to humans. The way of viral shedding depends on the respective tissue where the virus replicates: e.g. replication in kidneys and shedding via urine or replication in intestine organs and shedding via feces [62].

Transmission of viruses via saliva would be possible from bites when catching and handling the bats. Urine is constantly shed by the bats in the Wavul Galge cave, and the intake of these aerosols containing viral particles may lead to virus transmission when entering the cave without any protective equipment [63]. In this context, it would also be of interest in the future to investigate the virome of the other bat species inhabiting the cave and to conclude whether they are susceptible for the same viruses as well.

Feces is also constantly shed by the bats in their roosting cave. Although the fecal–oral transmission route is rather unlikely, local people are in close contact to bats when collecting bat guano to use as organic fertilizer [64]. Especially in rural areas like those around the Wavul Galge cave, the use of bat guano in agriculture is common and represents a potential transmission risk to the farmers. Although our study results revealed virus species of rather low zoonotic potential, this does not exclude the seasonal presence of potentially pathogenic agents. A special awareness regarding possible transmissions should be raised. Concurrently, the fear of zoonotic viruses in bat hosts should not justify their eradication. On the contrary, the natural habitat of the bat population in the cave should be recognized and respected.

In summary, the virome composition of different sample types obtained from *M. fuliginosus* bats in Sri Lanka was analyzed for the first time. Recent DNA barcoding and morphological studies on this species suggest that the *Miniopterus* bats inhabiting the island of Sri Lanka are in fact a new species of bat, not described hitherto and named as *Miniopterus phillipsi* [65]. Based on these findings, the results from our work would represent the first virome analysis for this newly described bat species.

## Acknowledgement

The authors are grateful to the MF2 unit at Robert Koch Institute for the sequencing of the samples and to Ursula Erikli for copy-editing. Further we thank the Department of Wildlife Conservation, Sri Lanka for granting the research permit.

