## Supplemental Material for "*Comparative virome analysis of individual shedding routes of* Miniopterus fuliginosus *bats inhabiting the Wavul Galge Cave, Sri Lanka*"

### Supplementary Table

Table ST1: Overview of the sample pools for NGS analysis, specifying included samples, total number of obtained reads after trimming and total reads that were assigned to viruses in MEGAN.

| Sample type | Pool | Included samples (Bat no.) | Total reads after trimming | Total reads assigned to viruses (MEGAN) |
|---|---|---|---|---|
| Oral swabs | O1 | 85, 87, 88, 89, 91, 94, 95, 96, 98, 100 | 4,906,956 | 3,005 |
| | O2 | 101, 103, 104, 106, 107, 108, 109, 110, 111, 113 | 5,381,629 | 1,757 |
| | O3 | 114, 115, 116, 117, 118, 119, 120, 121, 122, 123 | 5,416,413 | 3,317 |
| | O4 | 124, 125, 126, 127, 128, 129, 130, 131, 132, 133 | 4,568,104 | 1,732 |
| | O5 | 134, 135, 136, 137, 138, 139, 140, 142, 143, 144 | 4,609,884 | 1,776 |
| | O6 | 145, 146, 147, 148, 149, 150, 152, 153, 154, 156 | 6,206,508 | 2,584 |
| | O7 | 157, 158, 159, 160, 161, 162, 163, 164, 165, 166 | 4,984,406 | 2,731 |
| | O8 | 167, 168, 169, 170, 171, 172, 173, 174, 175, 176 | 4,570,809 | 1,153 |
| | O9 | 177, 178, 179, 180, 181, 182, 183, 184, 185, 186 | 3,241,858 | 2,412 |
| | O10 | 187, 188, 189, 190, 191, 192, 193, 194, 195, 196 | 4,515,414 | 1,629 |
| | O11 | 197, 198, 199, 200, 201, 202, 203, 204, 205, 206 | 3,963,803 | 3,142 |
| | O12 | 207, 208, 209, 210, 211, 212, 213, 214, 215, 216 | 3,794,828 | 1,578 |
| | O13 | 217, 218, 219, 220, 221, 222, 223, 224, 225, 226 | 4,025,694 | 1,881 |
| | O14 | 227, 228, 229, 230, 231, 232, 233, 234, 235, 236 | 4,549,502 | 2,165 |
| | O15 | 237, 238, 239, 240, 241, 242, 243, 244, 245, 246 | 3,726,854 | 2,247 |
| | O16 | 247, 248, 249, 250, 251, 252, 253, 254, 255, 256 | 5,625,662 | 1,465 |
| | O17 | 257, 258, 259, 260, 261, 262, 263, 264, 265, 266 | 8,428,422 | 4,434 |
| | O18 | 267, 268, 270, 271, 272, 273, 274, 275, 276 | 6,145,989 | 1,769 |
| | O19 | 277, 278, 279, 280, 281, 282, 283, 284 | 3,905,991 | 2,243 |
| Urine swabs | U2 | 87, 95, 96, 100, 101, 103, 104, 106, 111, 113 | 5,772,514 | 19,022 |
| | U3 | 114, 116, 227, 118, 119, 120, 132, 134, 135 | 4,231,010 | 99,135 |
| | U4 | 136, 137, 139, 143, 146, 147, 148, 155, 159, 160 | 3,626,749 | 80,589 |
| | U5 | 156, 166, 167, 168, 169, 170, 171, 172, 175, 176 | 5,557,808 | 113,844 |
| | U6 | 177, 178, 189, 181, 183, 184, 186 | 6,366,695 | 133,523 |
| | U7 | 187, 189, 190, 191, 193, 195, 196 | 9,849,006 | 209,126 |
| | U8 | 197, 198, 199, 201, 202, 203, 204, 205, 206 | 9,274,541 | 200,733 |
| | U9 | 207, 209, 217, 218, 225 | 8,084,957 | 252,039 |
| | U10 | 227, 229, 230, 232, 234, 236 | 4,264,597 | 90,724 |
| | U11 | 239, 240, 241, 242, 244, 246 | 3,652,926 | 76,205 |
| | U12 | 247, 248, 249, 253, 254, 255, 256 | 4,937,283 | 11,959 |
| | U13 | 258, 260, 261, 262, 263, 264, 265, 266 | 3,537,353 | 71,699 |
| | U14 | 267, 268, 270, 271, 272, 273, 278, 281, 284 | 922,687 | 17,963 |
| Feces | F1 | 91, 94, 100, 101, 113, 119, 123, 125, 127, 128 | 2,847,745 | 4,079 |
| | F2 | 131, 134, 142, 143, 147, 148, 149, 150, 153, 155 | 2,937,757 | 2,532 |
| | F3 | 156, 161, 173, 179, 180, 182, 183, 186, 189, 192 | 3,454,655 | 15,268 |
| | F4 | 195, 200, 202, 203, 204, 208, 209, 213, 215 | 2,404,303 | 3,173 |
| | F5 | 218, 220, 221, 222, 226, 227, 228, 230, 233, 234 | 2,921,352 | 2,426 |
| | F6 | 235, 236, 237, 238, 239, 241, 242, 243, 244, 245 | 1,497,502 | 882 |
| | F7 | 246, 247, 249, 250, 251, 252, 254, 256, 257, 258, 259 | 2,564,216 | 749 |

| | | | | |
|---|---|---|---|---|
| | F8 | 261, 263, 268, 270, 279, 280, 282, 283 | 2,993,081 | 898 |

#### Supplementary Figures

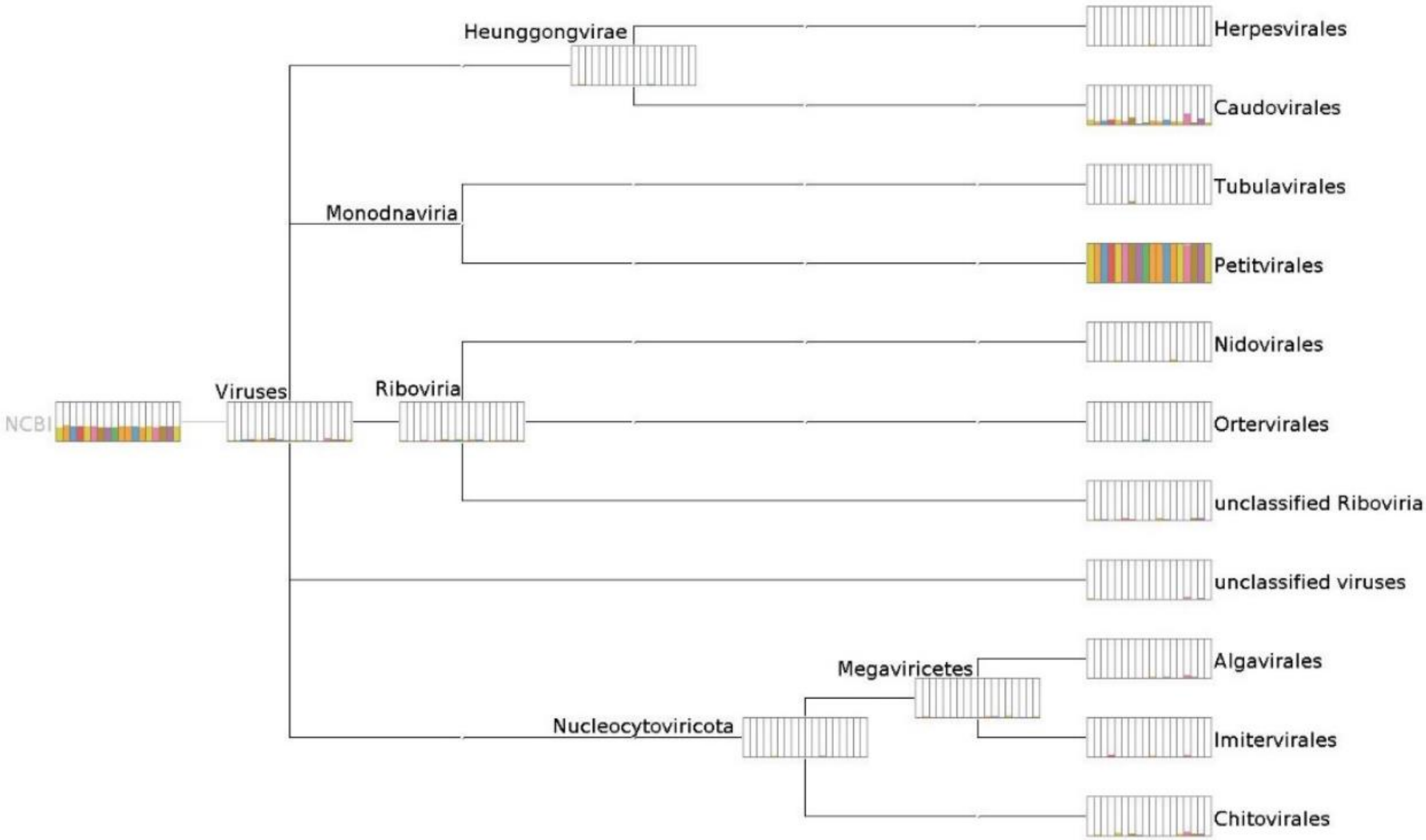


*Figure SF1: Normalized comparison of viral hits obtained after mNGS from OS pools OS2.1 – OS2.19, analyzed with diamond BLASTx algorithm and visualized in MEGAN software.*

### Supplementary Figures

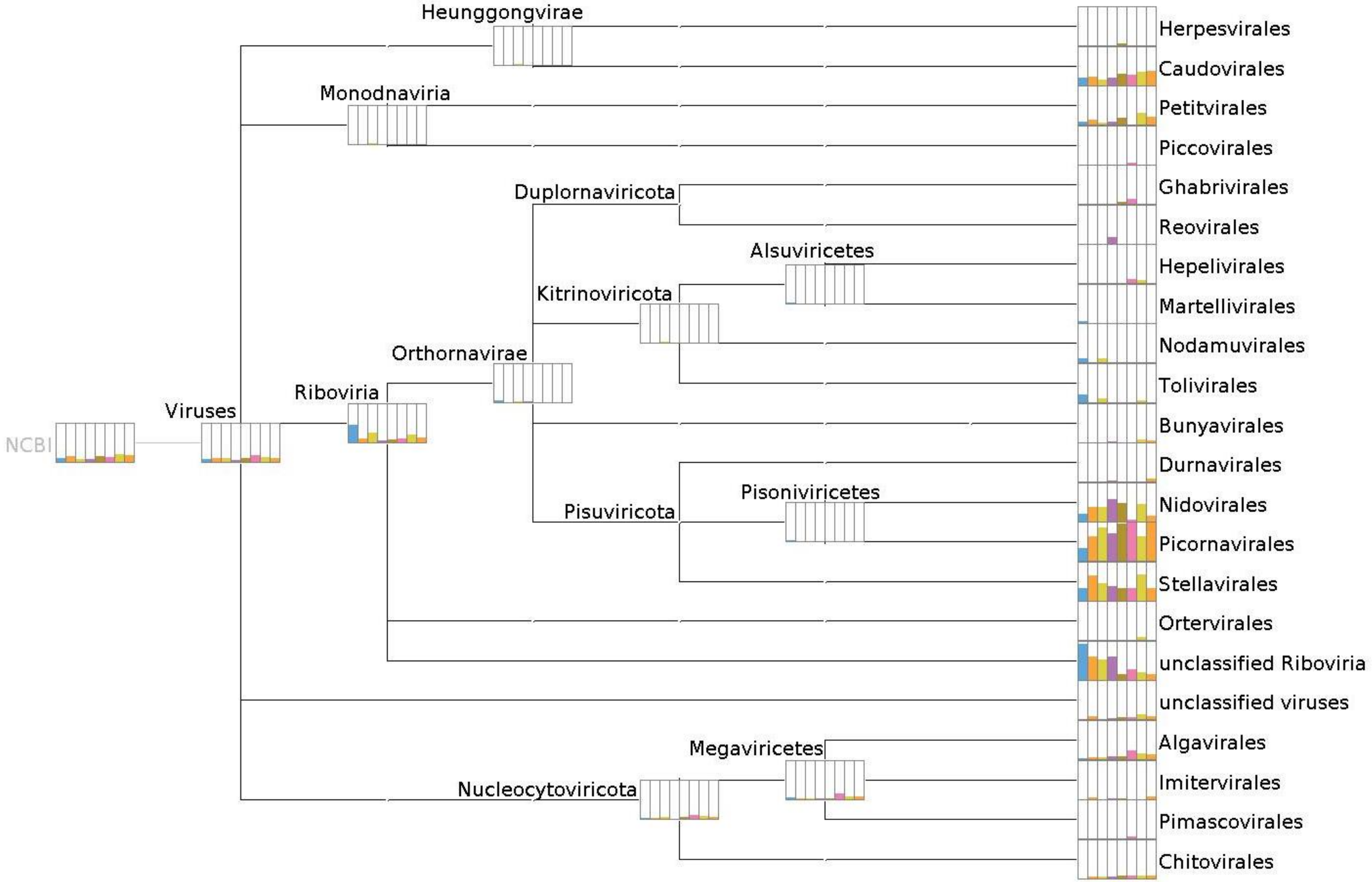


*Figure SF2: Normalized comparison of viral hits obtained after mNGS from F pools F2.1 – F2.8, analyzed with diamond BLASTx algorithm and visualized in MEGAN software.*

#### Supplementary Figures

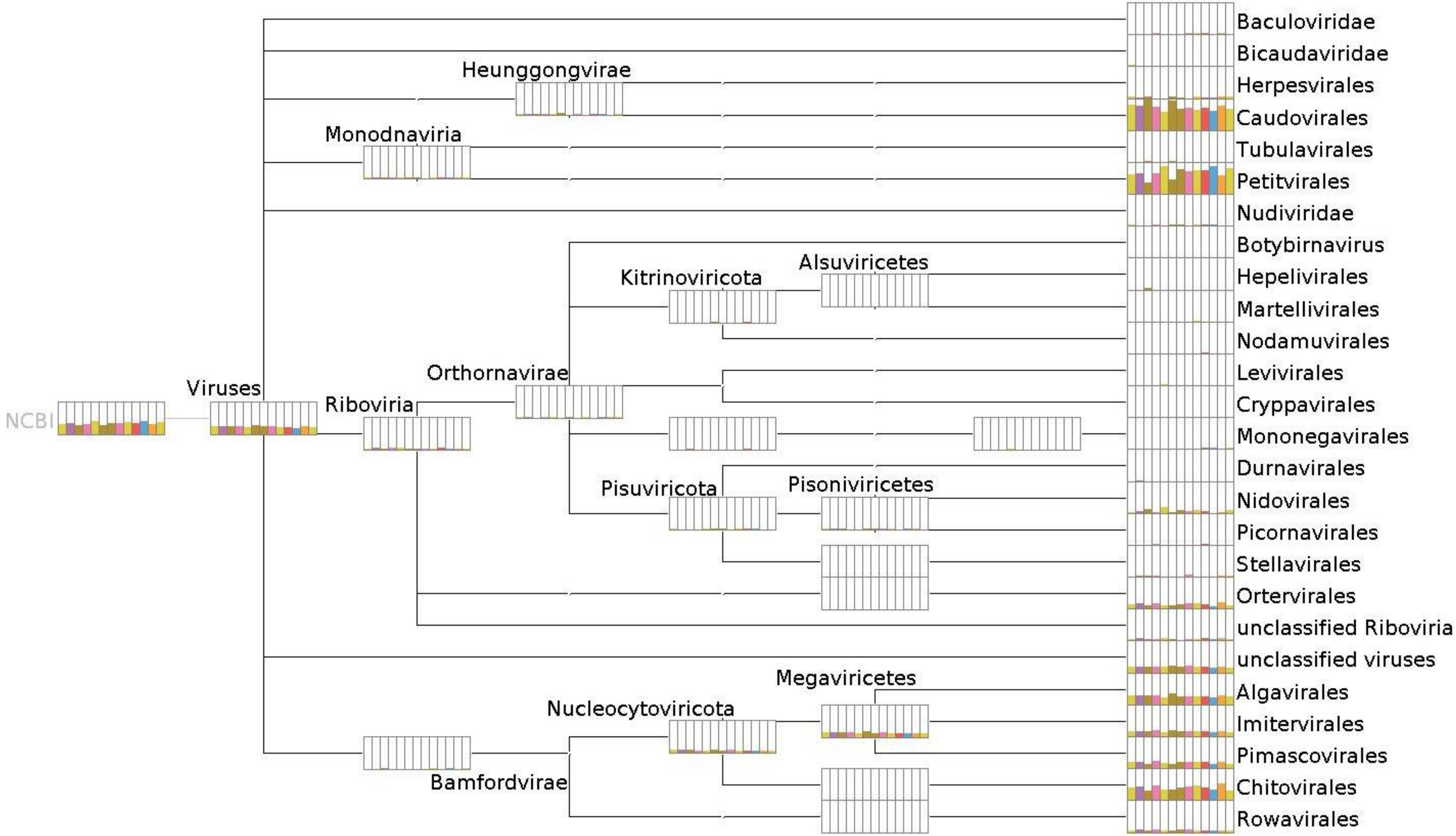


47 *Figure SF3: Normalized comparison of viral hits obtained after mNGS from US pools US2.1 – US2.14, analyzed with diamond BLASTx algorithm*
48 *and visualized in MEGAN software.*
